# Identifying the components of the *Shewanella* phage LambdaSo lysis system

**DOI:** 10.1101/2024.01.23.576932

**Authors:** Svenja Thöneböhn, Dorian Fischer, Vanessa Kreiling, Alina Kemmler, Isabella Oberheim, Fabian Hager, Nicole E Schmid, Kai M Thormann

**Affiliations:** Institute of Microbiology and Molecular Biology, Justus-Liebig-Universität Gießen, Gießen, Germany

**Keywords:** Prophage, holin, SAR-endonuclease, spanin

## Abstract

Phage-induced lysis of Gram-negative bacterial hosts usually requires a set of phage lysis proteins, a holin, an endopeptidase and a spanin system, to disrupt each of the three cell envelope layers. Genome annotations and previous studies identified a gene region in the *Shewanella oneidensis* prophage LambdaSo, which comprises potential holin- and endolysin-encoding genes but lacks an obvious spanin system. By a combination of candidate approaches, mutant screening, characterization and microscopy we found that LambdaSo uses a pinholin/signal-anchor-release (SAR) endolysin system to induce proton-leakage and degradation of the cell wall. Between the corresponding genes we found that two extensively nested open reading frames encode a two-component spanin module Rz/Rz1. Unexpectedly, we identified another factor strictly required for LambdaSo-induced cell lysis, the phage protein Lcc6. Lcc6 is a transmembrane protein of 65 amino acid residues with hitherto unknown function, which acts at the level of holin in the cytoplasmic membrane to allow endolysin release. Thus, LambdaSo-mediated cell lysis requires at least four protein factors (pinholin, SAR-endolysin, spanin, Lcc6). The findings further extend the known repertoire of phage proteins involved in host lysis and phage egress.

**Significance:** For the release of the assembled virions, phages have to breach the cell envelope. For Gram-negatives, this requires the disruption of three layers, the outer and inner membrane and the cell wall. In most cases, the lysis systems of phages infecting Gram-negatives comprises holins to disrupt or depolarize the membrane, thereby releasing or activating endolysins, which then degrade the cell wall. This, in turn, allows the spanins to become active and fuse outer and inner membrane, completing cell envelope disruption and allowing phage egress. Here we show that the presence of these three components may not be sufficient to allow cell lysis, implicating that also in known phages further factors may be required.

## Introduction

After completion of the infection cycle, many phages lyse their bacterial host cell to release their progeny into the environment. In Gram-negative bacteria, three barriers have to be breached: the cytoplasmic membrane (CM), the peptidoglycan of the cell wall (PG) and the outer membrane (OM) (reviewed in (1–3)). Lysis is initiated by holins (4), which are transmembrane proteins that, upon production, accumulate in the CM where they arrange to form non-specific lesions sufficiently large for the passage of phage-produced endolysins once phage assembly is complete (5). These endolysins are peptidoglycan-degrading enzymes, which, once released into the periplasm, perform the second step, the breakdown of the cell wall (2, 3). Alternatively, some phages possess non-canonical holins, so-called pinholins, in concert with signal-anchor-release (SAR) endolysins (6, 7). The latter are immediately transported into the periplasmic space after production but remain inactive by being tethered to the CM. Accumulation of the pinholins leads to formation of small transmembrane channels (7, 8), which results in breakdown of the proton motive force and the release of the SAR endolysins from the CM, upon which degradation of the cell wall proceeds. The third and final step of phage lysis is performed by spanins, which can occur in at least two versions as a unimolecular or a two-component spanin system (2, 3). In the more common two-component spanin systems, one component (the i-spanin) is integrated into the CM by a transmembrane region, while the second component (the o-spanin) is a lipoprotein, which is tightly anchored in the OM. Both components interact in the periplasm. According to the current membrane fusion model, the peptidoglycan layer prevents lateral diffusion of spanin units. Only upon hydrolysis of the cell-wall peptidoglycan by endopeptidase activity the spanin units can oligomerize and convert into a postulated hairpin conformation, which then mediates fusion of CM and OM, completion of cell lysis and free passage of the newly formed phage particles (2, 9–12).

However, for host cell lysis phages may also employ systems different from the canonical lysis components explained above. Some phages, such as *E. coli* phage Lambda (13), also produce antiholins, which can regulate timing of lysis initiation (14, 15). Some phages lack holins, such as *Escherichia coli* phage Mu, which instead uses a so-called releasin, a CM-tethered protein, to sever the association of the corresponding SAR endolysin with the membrane (16). Other phages lysing Gram-negatives do not possess apparent spanins. They may be replaced by alternative systems, such as in *E. coli* phage phiKT, where disruption of the OM is taken over by a cationic membrane-associated protein (17).

The facultative gammaproteobacterium *Shewanella oneidensis* harbors the Lambda-like prophage LambdaSo. Previous studies showed that induction of the LambdaSo lysogenic cycle occurs during early surface attachment and that subsequent phage-induced cell lysis is an important source of extracelluar DNA, which in turn is required for biofilm formation and to maintain the biofilm’s structural integrity (18–20). Accordingly, deletion of the annotated LambdaSo prophage lysis gene cluster (21) resulted in cells unable to release intracellular beta-galactosidase and to form normal biofilms, strongly indicating that one or more genes in this cluster are, in fact, required for cell disruption (20). While genes annotated as holin and potential endolysin are present in this cluster, spanin-encoding genes were not identified. This prompted us to revisit, identify and functionally characterize the components of the LambdaSo lysis system. We found that for phage egress LambdaSo relies on a system consisting of a (so far falsely annotated) pinholin together with a SAR-endolysin and a two-component spanin system. In addition, we found that a novel membrane protein factor is strictly required for LambdaSo-induced cell lysis. This small transmembrane protein likely acts at the level of pinholin at the early step of lysis.

## Results

The LambdaSo gene region SO_2966 - SO_2974 has previously been shown to be essential for lysing the host cell (20). This region comprises genes that were originally annotated as a putative holin (SO_2969) and lysozyme (SO_2973), although their function had not been experimentally proven (**Figure 1**). In a first set of experiments we therefore aimed at validating the function of these two lysis components. To this end, we used an *S. oneidensis* background strain in which two other prophages in *S. oneidensis*, MuSo1 and MuSo2, were deleted from the strain to avoid interference with LambdaSo-derived phenotypes. In the following, this strain will be referred to as the wild type for simplicity. To firstly determine the time frame of phage release after LambdaSo induction, we performed a one-step growth curve. After addition of mytomycin C to trigger the cellular SOS response and LambdaSo activity, phage release started after about 100 min and about 250 infectious particles were released per cell (**Supplementary Figure 1**).

**Figure 1:**
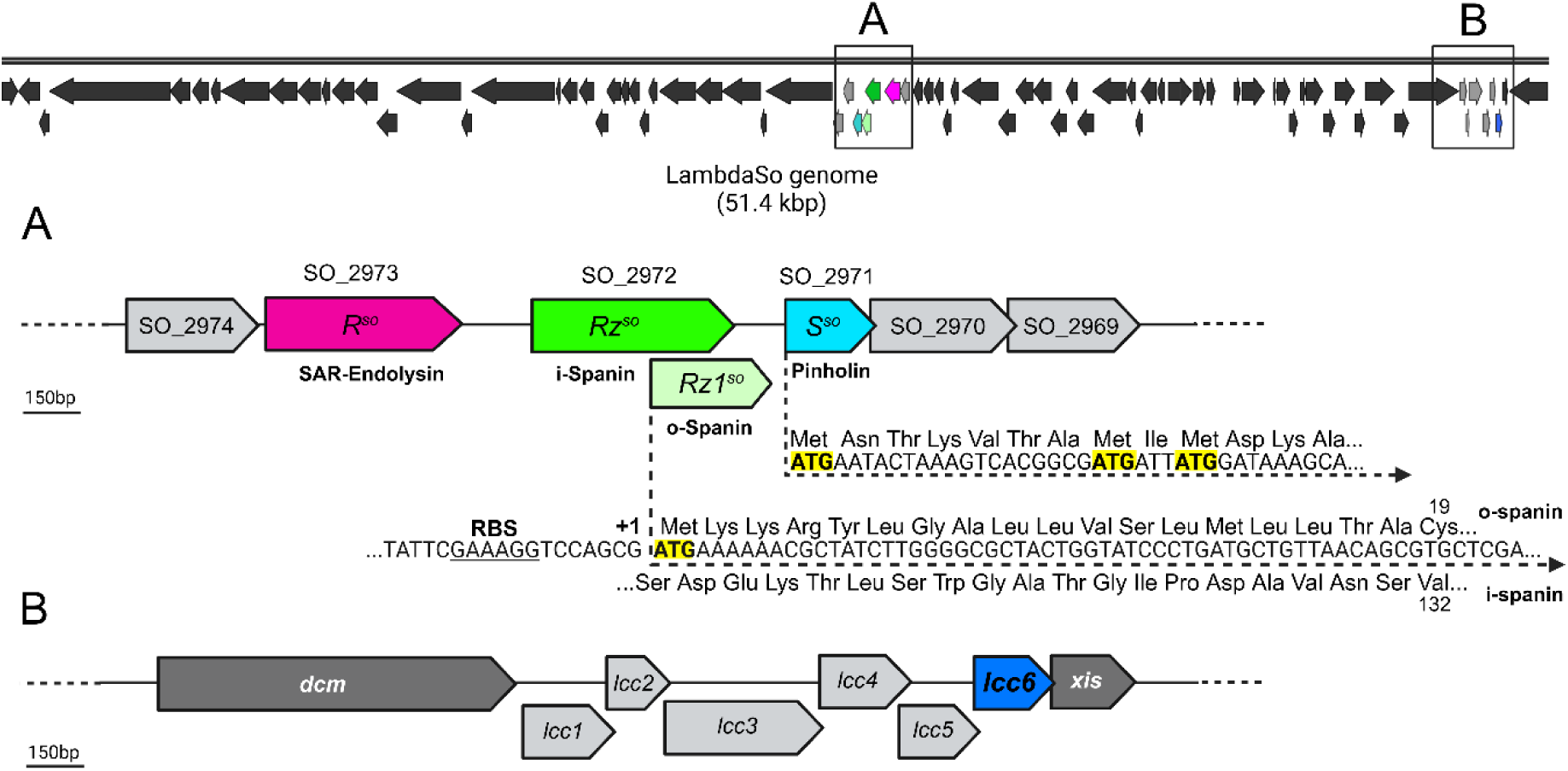
Genetic organization of *Shewanella* prophage LambdaSO. The upper panel displays the full annotated genome of LambdaSO. The predicted open reading frames within the phage genome are represented as arrows indicating their transcriptional direction. The boxes labeled A and B show the position of the gene clusters involved in phage-induced cell lysis, which are displayed in the middle and lower panels in more detail. The deduced gene products are color-coded according to their predicted function. **A.** Genetic organization of the predicted lysis cluster (SO_2971 to SO_2973). Genes encoding proteins directly involved in cell lysis are indicated as coloured arrows. The the predicted translation start of Rz1^So^ within the +1 frame of the gene encoding Rz^So^ is highlighted in yellow, the upstream ribosome binding site is also indicated. The three alternative start sites of the pinholin-encoding gene S^So^ are accordingly highlighted in yellow. Bioinformatic predicitons are based on NCBI BLASTP (National Library of Medicine) and PHAST analyses (52). **B.** Gene organization of so-called LambdaSo cluster C (*lcc1* (SO_4794) to *lcc6* (SO_4975)). Predicted genes are indicated as arrows. *lcc6*, which is part of the lysis machinery, is colored in blue, the other *lcc* genes are colored in light grey, the neighboring genes are shown in dark grey. Bioinformatic predictions are based on NCBI BLASTP (National Library of Medicine).

### Identification of SO_2971 as LambdaSo holin

In the original annotation of the *S. oneidensis* genome, the gene product SO_2969 was identified as the putative holin. Notably, this protein (109 aa) does not exhibit any homologies to known holins and also lacks transmembrane domains. Instead, SO_2969 has high homologies to so-called HNH proteins, which were shown to be part of the phage terminase DNA packaging system in numerous phages (22). Accordingly, the gene is located directly upstream of that encoding the small subunit of the LambdaSo terminase (SO_2968). We therefore concluded that the original annotation is incorrect. Homology searches identified another holin candidate (SO_2971) encoded two genes upstream. The deduced gene product (89 aa, **Supplementary Figure 2**) exhibits clear homology with respect to sequence and membrane topology to predicted holins of the HP1 family from other phages (4). The homologies thus suggested a function of SO_2971 as a so-called pinholin with two transmembrane domains. In addition, the open reading frame harbors two additional alternative translational start sites six and eight amino-acids positions downstream of the annotated start codon (**Figure 1**). These may lead to the production of slightly shorter holin variants, suggesting that LambdaSo possesses a holin/antiholin system as described for the pinholin system of *E. coli* phage ɸ21 (23). We therefore speculated that SO_2971, hereafter referred to as LambdaSo holin S (S*^So^*), is the sought-after protein to initiate host lysis by depolarizing the cytoplasmic membrane.

To determine the function of the putative holin, the corresponding gene was deleted from the phage chromosome (Δholin; ΔS^So^), which would leave the phage without holin/antiholin system. Upon Lambda induction the wild-type cells as well as those of the Δholin strain exhibited elongation, however, as opposed to wild-type cells, cells of the Δholin strain failed to lyse and no plaque-forming particles are found in the culture supernatant (**Figure 2; Supplementary Figure 3**). Reintegration of the gene restores normal lysis (**Supplementary Figure 4**). In addition, we determined a potential holin-induced membrane depolarization via DiBAC staining. Although depolarization is expected to occur very shortly before cell lysis, which could not be synchronized within the culture in such a short time frame, a significant difference in polarization could be determined between wild type- and Δholin cells (**Figure 3**). Based on these results we concluded that SO_2971 serves as a pinholin in the LambdaSo lysis system.

**Figure 2:**
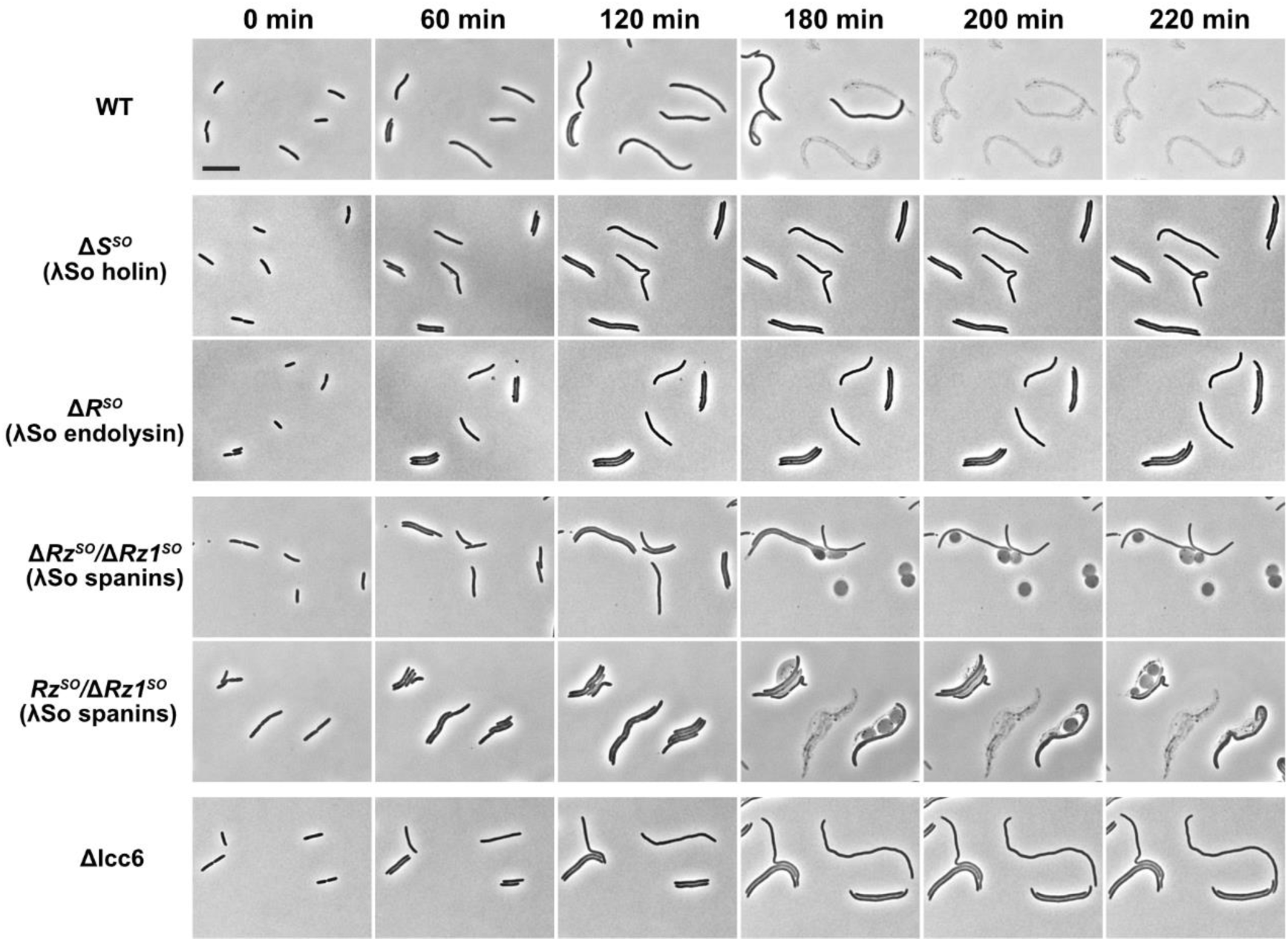
LambdaSo-induced lysis of *S. oneidensis* MR-1 requires a pinholin, SAR endolysin, spanin complex and the small transmembrane protein Lcc6. The micrographs represent timelapse series of *S. oneidensis* strains in which the genes encoding for the pinholin (*S^SO^*), SAR endolysin (*R^SO^*), spanin complex (*Rz^SO^/Rz1^SO^*) and Lcc6 (*lcc6*) were deleted from the λSo genome as indicated. The time points subsequent to LambdaSo induction by addition of mitomycin C (10 µg/ml) are indicated above. The scale bar represents 5 µm. The corresponding complementation strains are shown in **Supplementary Figure S4**.

**Figure 3:**
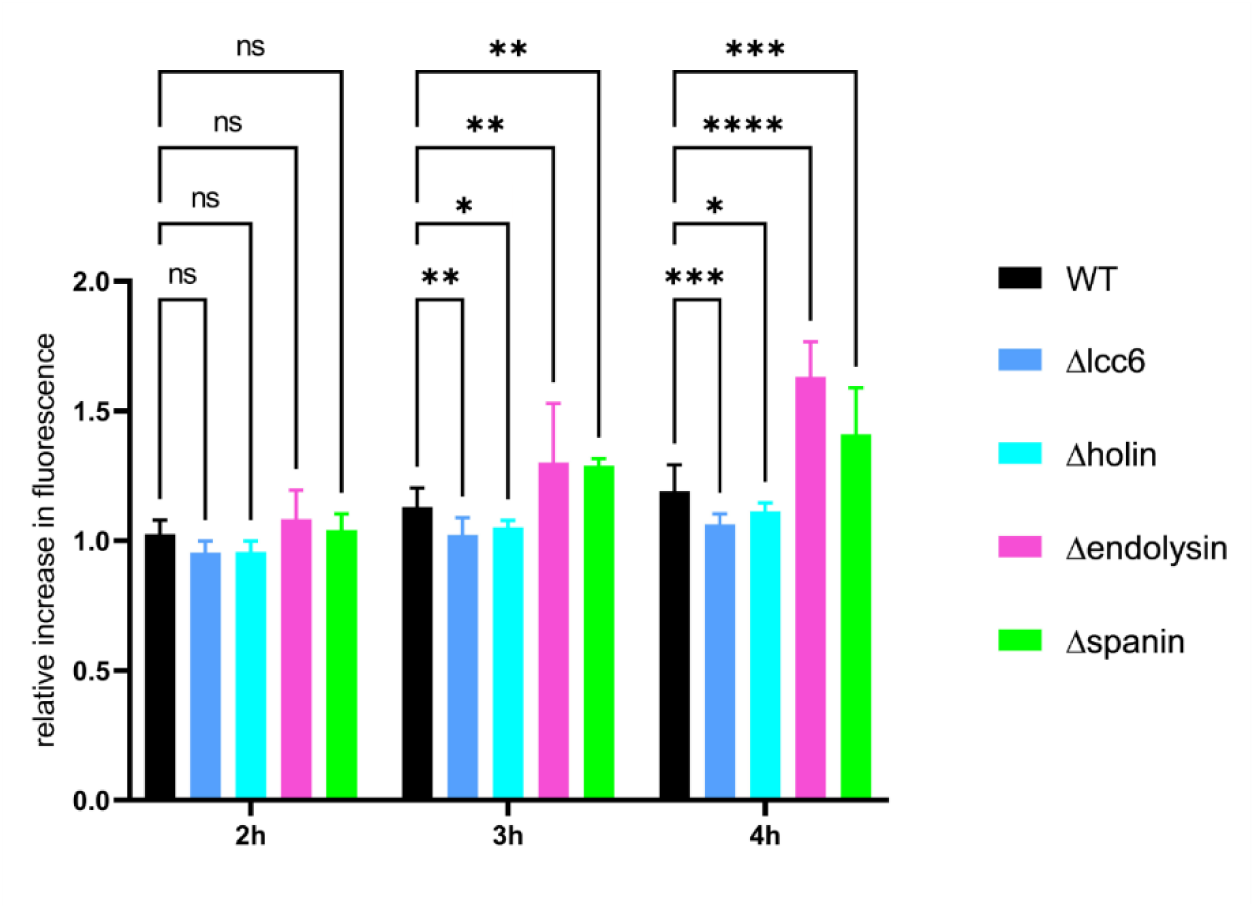
Membrane depolarization assay on LambdaSo mutants using DiBAC4(3). The cells were grown to exponential prior to LambdaSo induction by mitomycin C phase and stained with DiBAC4(3) to visualize depolarization after the indicated time points. Statistical significance is indicated by the p-value. NS = not significant, * p-value = P ≤ 0.05, ** p-value ≤ 0.01, *** p-value ≤ 0.001, **** p-value ≤ 0.0001

### SO_2973 as the LambdaSO endolysin

SO_2973 was originally annotated as lysozyme and was thus the obvious candidate to be the lysis component required for LambdaSo to breach the peptidoglycan after holin-induced pore formation in the CM. SO_2973 consists of 170 aa and is predicted to possess an N-terminal Sec signal peptide but is lacking a clear proteolytic cleavage site following the signal peptide (structure prediction see: **Supplementary Figure 2**). Thus, the protein likely belongs to the group of signal-anchor-release (SAR) endolysins, which are exported into the periplasmic space but remain tethered to the cytoplasmic membrane by their N-terminal non-processed signal peptide until being released through membrane depolymerization by pinholin-mediated pore formation (6, 24–26).

To determine the function of SO_2973 for LambdaSo-induced cell lysis we created an in-frame deletion in the corresponding gene. As observed for the deletion of the holin, the cells elongated after phage induction, but the cells failed to lyse (**Figure 2**). Accordingly, no plaque-forming phage particles were found in the supernatant (**Supplementary Figure 3**). Phage-induced lysis was restored by reintegration of the gene (**Supplementary Figure 4**). Notably, as opposed to the holin mutant, significant depolarization of the membrane occurred in SO_2973 mutants (now *R^So^*) (**Figure 3**), indicating that the first step of lysis, executed by the holins, is carried out but the barriers of the cell wall and the outer membrane remain intact.

### Identification of phage protein SO_4795 (Lcc6) as a factor involved in cell lysis

While holin and endolysin were readily identified, a candidate gene product showing homologies to known spanins was neither annotated nor could be identified by genome searches. It has been shown that some phages infecting and lysing Gram-negative hosts do not possess spanins, and a recent study identified a cationic antimicrobial peptide that alternatively assists in disrupting the outer membrane (17). We assumed that this may be similarly true for LambdaSo and therefore explored the role of small open reading frames within in the LambdaSo gene region for a potential role in cell lysis.

As the vast majority of phages, LambdaSo harbors a number of genes with no annotated functions. One predicted gene cluster consists of six genes, hereafter referred to as *L*ambda *c*luster *C* (***lcc***) 1 to 6, and is located at the far end of the LambdaSo genome directly upstream of the *xis* gene (**Figure 1**). The *lcc* genes encode rather small proteins with unknown functions, two of which are predicted to possess a transmembrane region (*lcc4* and *lcc6*). Deletion of the *lcc* gene cluster from the genome resulted in a drastic decrease in the production of active phage particles in the supernatant, which would be an expected phenotype for a lysis-deficient mutant of LambdaSo (**Supplementary Figure 5**). We performed gene deletion of genes *lcc2* to *lcc6*, a single deletion of *lcc1* could so far not been obtained. Each of the resultant strains was tested with respect to its ability to produce infectious particles in the supernatant. Among these genes, a deletion of *lcc6* phenocopied the deletion of the full cluster, which could be complemented by reintegration of the gene into its original locus (**Supplementary Figure 5**). Loss of *lcc2*, *lcc3*, *lcc4* and *lcc5* had no significant effect on LambdaSo propagation under the conditions tested.

*lcc6* is predicted to encode a protein of 65 aa residues with an N-terminal transmembrane region (aa 5-22). Plasmid-borne production of Lcc6 had no apparent effect on cell morphology (**Supplementary Figure 6**). To determine the effect of Lcc6 on LambdaSo production, the corresponding mutant was observed microscopically after induction of the LambdaSo lytic cycle by addition of mitomycin C. When lacking Lcc6, the cells elongated after phage induction, but failed to lyse (**Figure 2**) in a way that is highly reminiscent to the phenotype of holin or endolysin mutants. In addition, the Lcc6 mutant exhibited a limitation in membrane polarization and thus resembled a holin deletion phenotype rather than that of an endolysin deletion (**Figure 3**). The data indicated that Lcc6 is strictly required for phage-induced cell lysis and suggested that the protein acts at the level of (or together with) the holin and is not directly involved in disintegrating the outer membrane.

### Two nested genes (SO_2972a and SO_2972b) encode the LambdaSo spanin system

In a complementary approach to that of addressing the role of small phage proteins in outer membrane disruption to allow phage egress, we aimed at the identification of potential spanins by a candidate approach. In *E. coli* phage Lambda and other phages, the genes encoding the necessary factors for cell lysis, holin, endolysin and spanin (*S, R, Rz)* are directly located next to each other within the phage genome. We therefore hypothesized that the LambdaSo spanin might be similarly encoded in proximity to the holin- and lysozyme-encoding genes (see **Figure 1**). The gene upstream of the SAR-endolysin *So*^R^ encoding gene, SO_2974, is predicted to encode a pyridoxal phosphate-dependent protein with no apparent transmembrane region to suggest a function as spanin. SO_2970, encoded downstream of the holin S, is a protein of unknown function also lacking any apparent transmembrane region. In contrast, SO_2972, located between *R^So^* and *S^So^*, is a protein of 174 aa predicted to possess an N-terminal transmembrane region (aa 9-31) followed by a long alphahelical stretch (aa 32-110) (**Supplementary Figure 2**). The predicted topology of the protein would fit well to the cyto- and periplasmic component of a two-component spanin system (see (2, 3)). In *E. coli* phage Lambda, but also in other phages (27), the second spanin component, Rz1, a lipoprotein making contact with the outer membrane, is encoded as a second open reading frame within the *Rz* gene (9, 28). Genetic analysis revealed that, similarly, a second potential open reading frame (SO_2972b) starts within SO_2972 at the +1 frame starting at position 340. A potential Shine-Dalgarno sequence (GAAAGG) is located 7 bp upstream of the predicted start codon. The open reading frame extends into the gene region between SO_2972 and SO_2971, encoding a protein of 95 aa with a lipoprotein signal peptide to be cleaved between position 18 and 19 (**Supplementary Figure 2**), leaving an N-terminal cysteine residue required to attach the outer membrane lipid anchor (29).

To determine the role of SO_2972a (*Rz^So^*) and SO_2972b (*Rz1^So^*) as a potential two-component spanin system, SO_2972a was deleted from the genome, which also inactivated the putative overlapping gene SO_2972b. To specifically target SO_2972b, we introduced a GTG to GTC base substitution within SO_2972a (V132) which does not change the Rz protein but results in a Cys19Ser substitution in SO_2972b, removing the cysteine residue thought to serve as the lipid attachment site **(Supplementary Figure 2**).

The resultant mutants lacking *Rz^So^*/*Rz1^So^*or *Rz1^So^* were then observed by microscopy after LambdaSo induction. Cell elongation proceeded like in wild-type cells, however, a number of the cells converted into spheres instead of lysis in both mutants (**Figure 2**). Previous studies have shown that spanins are particularly important for phage egress when the cells are growing under conditions where the outer membrane is stabilized, e.g., in the presence of Mg^2+^ ions (30). When grown in planktonic shaking cultures without addition of Mg^2+^, infectious plaque-forming phage particles were found in the medium supernatant, albeit at a number of 3 to 4 orders of magnitudes less than wild-type phages. However, under membrane-stabilizing conditions, no plaques were produced by *Rz^So^*/*Rz1^So^*mutants (**Figure 4; Supplementary Figure 3**). Accordingly, also during time-lapse microscopy spheres formed by *Rz^So^*/*Rz1^So^* mutants were more stable, which allowed to microscopically follow sphere formation. These morphology changes commonly started at one cell tip and then proceeded through the elongated cell at a time frame of about 1 min (**Figure 4**). Taken together, the results strongly indicate that Rz*^So^* and Rz1^So^ constitute a two-component spanin system of phage LambdaSo.

**Figure 4:**
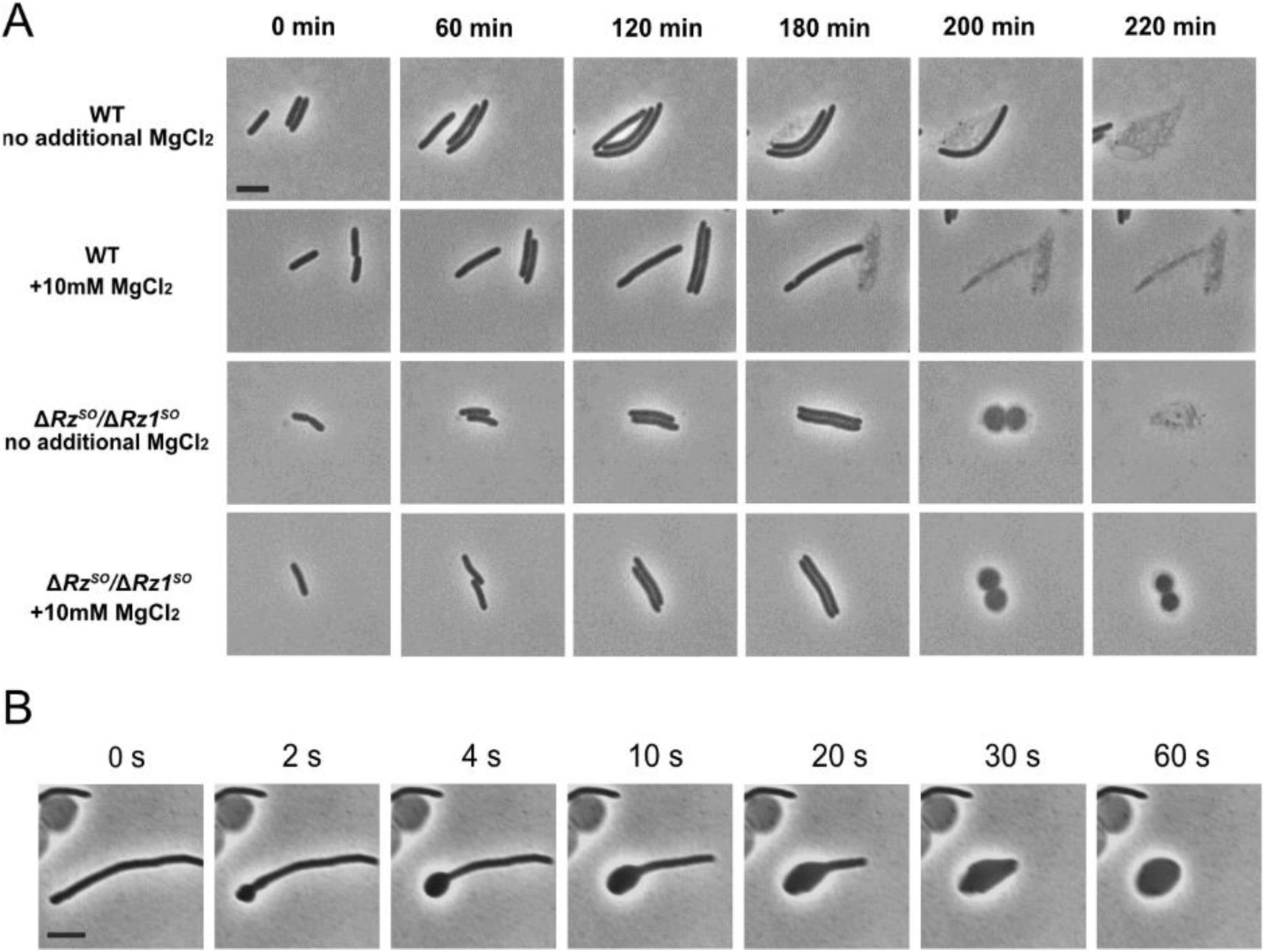
The spherical cell morphology of spanin-lacking mutants is stabilized by Mg^2+^ and initiates at the cell pole. **A.** Displayed are micrographs of a timelapse series of *S. oneidensis* wild-type and Δspanin (Δ*Rz^SO^/*Δ*Rz1^SO^)* cells treated with and without 10 mM MgCl_2_ supsequent to phage induction of the wild type (WT) and the spanin-lacking LambdaSo prophages using mitomycin C (10 µg/ml). The time points are indicated above; time point zero is defined as the start of induction. The scale bar represents 2 µm. **B.** Morphology changes of cells lacking a spanin system after phage induction. Micrographs were taken after the time points indicated above. Full conversion into a sphere takes about 60 seconds. The scale bar represents 2 µm.

## Discussion

*Shewanella* phage LambdaSo is an active prophage in *S. oneidensis*, which has been shown to exert a pronounced effect on the physiology of its host, e. g., with respect to biofilm formation and repression of other prophages (18–20, 31). LambdaSo induction requires the RecA protein and occurs in the wake of the host’s SOS response (19, 20). We found that under our standard conditions, about 250 phages per cell are released and cell lysis starts after about 120 minutes after inducing the host’s SOS response. Here, we characterized the phage-induced lysis of *S. oneidensis* and identified the components required for LambdaSo egress, a (pin)holin, a SAR-endolysin and a two-component spanin system, and we found an additional novel phage factor involved in host lysis.

### The pinholin/SAR-endolysin of LambdaSo

The initial annotation of the *S. oneidensis* MR-1 sequence (21) designated a putative function to a number of LambdaSo phages and identified a gene region comprising a holin- and a lysozyme-encoding gene. Accordingly, a deletion of the gene region resulted in *S. oneidensis* mutants releasing less extracellular DNA during initital biofilm formation (20), supporting the role of the genes in this cluster in phage-induced cell lysis. While the initial annotation for the holin was incorrect, the correct holin-encoding gene, SO_2971 (now LambdaSo holin S, S*^So^*) was readily identified by homology comparisons of the gene products from neighboring genes. According to the homologies, the LambdaSo holin belongs to the group of pinholins, which, as opposed to canonical holins, harbor two (instead of three) transmembrane regions and only form small pores at the nanometer scale to disrupt the proton motive force (pmf) (7, 8). Pinholins usually occur together with a special class of endolysins, the SAR-endolysins, which already reside in the periplasmic space but are kept inactive by an N-terminal transmembrane domain that tethers the protein to the CM. Breakdown of the pmf releases and activates the SAR-endolysins (6, 7) and licenses initiation of the peptidoglycan degradation. Accordingly, the Lambda*So* endolysin (now LambdaSo endolysin R, R*^So^*) exhibits high homologies to other proteins of that group and sports the N-terminal signal/transmembrane-anchoring domain. Thus, we concluded that LambdaSo-induced pmf collapse and cell-wall degradation occurs via a pinholin/SAR-endolysin pair.

The pinholins of phage ɸ21 are the best-studied representatives of that holin type and are encoded by the gene S^21^. The gene has two translational start sites (23, 32), leading to the production of two pinholin isoforms, the full length S^21^71 pinholin and the slightly smaller S^21^68 variant that lacks three N-terminal residues. Both variants possess two TM regions, where the N-terminal TM1 serves as a negative regulator and TM2 acts as the hole-forming domain (8). The smaller variant S^21^68 is the lysis effector, as its TM1 is readily able to externalize from the membrane into the periplasm (33). This results in the assembly of the holin units into monoheptameric microchannels, which subsequently allow the passage of protons and membrane depolymerization. The slightly longer holin version, S^21^71, possesses an extra positively charged residue at the cytoplasmic site of the protein’s N-terminus, which impedes the externalization of TM1. By forming non-active membrane-embedded heterodimers with S^21^68, S^21^71 serves as an antiholin for the shorter active version. Ratio and accumulation of the two variants mediate proper timing of cell lysis (8, 34–36). The pinholin of LambdaSo, S*^So^*, has even two alternative translational starts, which could – in theory – result in the production of three isoforms, full-length S*^So^*89, S*^So^*82 and S*^So^*80. The S*^So^*89 version would, similarly as S^21^71, possess an additional positively charged amino acid, a lysine at position 4. We therefore hypothesize that S*^So^*89 similarly functions as an antiholin for one or both of the shorter isoforms. However, this remains to be studied in more detail.

### LambdaSo-induced lysis involves a two-component spanin system

While two of the LambdaSo main lysis components were readily identified, no spanin components turned up by homology searches. Known spanins consist of one or two protein components (reviewed in (2, 3)), but have in common that interaction with the OM is mediated via a lipid anchor. In Gram-negatives such proteins are characterized by a signal sequence followed by one or more cysteine residues to which the lipid anchor is coupled (29). LambdaSo harbors two genes located away from the lysis cluster (SO_2991 and SO_2938), which are predicted to encode potential lipoproteins. Notably, in our hands these genes could not be deleted from the LambdaSo chromosome; thus, their roles for the phage remain elusive so far. The LambdaSo spanin system comprising the i-spanin Rz*^So^* and the o-spanin Rz1*^So^* was then identified by a candidate approach concentrating on the genes located in the neighborhood of those encoding holin and the lysozyme. As previously shown for number of phages including *E. coli* Lambda (9, 28), the genes encoding Rz*^So^*and Rz1*^So^* are intimately nested. Rz1 starts in the 3‘-half of Rz and extends towards the gene encoding the pinholin S*^So^*, thereby bridging the gap that was originally annotated as non-coding region. Thus, the four genes are likely forming a continuous lysis cluster (**Figure 1**). Alphafold predictions show that Rz*^So^* has the shape of a long alphahelix with the N-terminus anchored in the CM, followed by a turn and a short second alphahelix, which is somewhat reminiscent to that of the paradigm two-component spanin of *E. coli* Lambda. In addition, Rz*^So^* possesses a residual stretch of residues without predicted folding (**Supplementary Figure 2**). Cysteine residues, which may assist in dimerizarion *in vivo* by disulfide bonding, as shown for *E. coli* Lambda (37), are present at positions 139 and 150.

With the signal sequence removed, the N-terminus of Rz1*^So^*starts with the leading cysteine for lipid fusion followed by a stretch without any predictable structure and a C-terminal alphahelix that is not present in the *E.coli* Lambda counterpart. An extensive accumulation of proline residues comparable to that found in Rz1*^Ec^* is absent, two cysteine residues potentially involved in homodimerization are present at position 47 and 74, as opposed to a single cysteine residue in *E. coli* Lambda Rz1. Thus, while the general mechanism of membrane fusion is likely conserved between the two systems, the domains and residues of LambdaSo required for spanin complex formation are different from those identified in *E. coli* Lambda (38, 39). The interaction between the two components is currently under investigation.

### Lcc6, a novel player in phage-induced lysis

Surprisingly, we identified another protein to be required for proper cell lysis and phage release. This protein, which we termed Lcc6, is encoded elsewhere on the LambdaSo phage genome as part of a cluster consisting of six small open reading frames, which is located close to the predicted *int*/*xis* genes almost at the border region of the phage genome. Lcc6 consists of a transmembrane domain followed by a stretch of 43 aa that is predicted to remain in the cytoplasm. The protein is essential for lysis as in the absence of the protein, phage release does not occur. On the other hand, overproduction of Lcc6 without simultaneous phage induction has no apparent effect on cell morphology or growth and therefore is not a sole lysis factor but likely acts in concert with other LambdaSo proteins. The absence of cell rounding after phage induction and the observed little change in membrane potential in *Δlcc6* mutants strongly suggests that Lcc6 functions at the level of pinholin pore assembly and SAR-endolysin release from the cytoplasmic membrane.

Chamblee et al (16) recently identified the long missing protein that allows the SAR-endolysin of *E. coli* phage Mu to be released and activated, as this phage lacks a holin system. Instead, this phage employs a membrane-localized DUF2730 protein, which was accordingly designated releasin. The mechanism by which releasin acts is not completely clear but it is proposed that the protein directly or indirectly neutralizes the positive charges at the cytoplasmic face of the endolysin’s transmembrane domain, e. g., by removal through endolytic cleavage or formation of a hydrophylic channel (16). Lcc6 does not exhibit notable homologies to the Mu releasin gp25, does not have a DUF2730 domain and is also shorter. In addition, in the absence of Lcc6 membrane depolarization is decreased compared to that of holin mutants, making a role of this protein in facilitating endolysin release rather unlikely and implicates a role at the level of the holin activity. As detailed above, LambdaSo may possess a complex pinholin set-up with up to three isoforms with at least one anti-pinholin. It may therefore be hypothesized that Lcc6 is involved in triggering pinholin assembly, e.g., by antiholin removal or directly facilitating formation of the pore complex. So far, we have no direct experimental evidence if and how such an Lcc6 activity may occur and how this may affect the timing of cell lysis. However, the finding of another protein factor in addition to a (anti-)holin/(SAR-)endolysin/spanin system required for cell lysis was unexpected and raises the question if more Lambda-like or other *Caudovirales* phages possess such Lcc6-like factors.

## Materials and Methods

### Cultivation of bacterial strains

All bacterial strains used in this study are summarized in **Supplementary Table 1**. *S. oneidensis* cells were grown in LB medium at room temperature or 30°C, cells of *E. coli* in LB medium at 37°C. Media for *E. coli* WM 3064 [2,6-diaminoheptanedioic acid (DAP)-auxotroph] were supplemented with DAP at a final concentration of 300 μM. Selective media were supplemented with 50 mg/ml Kanamycin, 10 mg/ml Chloramphenicol or 10 mM MgCL_2_.

### Induction and cultivation of phages

To derive LambdaSo from the respective *S. oneidensis* strain, the phages were induced either through addition of 10 µg/ml mitomycin C to an exponentially growing liquid culture of the corresponding strains or an incubation of the culture at 30°C at 40 rpm for ∼48 h. To prepare the phage lysate, the entire cell material was then pelleted in a centrifuge at 13,000 rpm for 15 min or 3500 rpm for 5min at room temperature, which was followed by a filtration of the supernatant using a 0.25 µm filter. To determine the plaque forming units (pfu) per ml of the phage lysate, a plaque assay was performed.

### Plasmid and strain constructions

Plasmids and corresponding oligonucleotides used in this study are summarized in **Supplementary Tables 1 and 2**. For DNA preparations, the appropriate enzymes (Fermentas, St. Leon-Rot, Germany) and kits (VWR International GmbH, Darmstadt, Germany) were used. Gene deletion and insertion strains were generated by sequential double homologous recombination using the suicide plasmid pNPTS-R6K (40). Briefly, substitution fragments were constructed by combining approximately 500-bp fragments of the up- and downstream regions of the designated gene. For gene deletions, a few codons (typically around six) of the corresponding reading frame were kept. Single or multiple nucleotide substitutions were done by inserting a modified gene fragment into the corresponding gene deletion. Plasmids were constructed using standard Gibson assembly protocols (41) and transferred into *Shewanella* cells by electroporation or conjugative mating with *E. coli* WM3064.

### Phage functionality / Lysis assay

To verify the ability of phage induced host lysis spot test analysis were performed. Phage lysates were isolated from the corresponding bacterial strains which were incubated for 48 h (30 °C) by centrifugation (3500 rpm, 5min). 1.6 ml of an exponential growing bacteria suspension lacking the λSo genome was inoculated in 30 ml 0.5 % soft-agar supplemented with or without 10 mM MgCl_2_ (30) and spread on LB plates. 3 μl of individual phage lysates were spotted onto the surface of the plates. The lysing properties of the phages were displayed by forming clear zones on the bacterial lawn.

### Microscopy

Samples for all microscopy images were harvested from exponential growing *S. oneidensis* cultures. Microscopy images were acquired by a DMI6000 B inverse microscope (Leica) equipped with a pco.edge sCMOS camera (PCO) and an HC PL APO 100x/1.40 oil PH3 phase contrast objective utilizing the VisiView software (Visitron Systems GmbH). Images were subsequently analyzed and edited using the ImageJ-based Fiji tool (42), PowerPoint 2019 (Microsoft Corporation) and Affinity Designer 1.7v (Serif).

### One-step growth experiment

To determine the latent period and burst size of LambdaSo, a one-step growth experiment was performed (43–45). Briefly, 20 ml of cell culture of *S. oneidensis* ΔLambdaSo ΔMuSo1 ΔMuSo2 was prepared in LB medium and incubated at RT. On the following day, 160 ml of LB medium was mixed with 1.6 ml of the grown culture and left to stand on the shaker at 32 °C up to an OD_600_ of ∼0.4. The entire preparation was then divided into 3 x 50 ml samples and centrifuged at 6000 rpm for 15 min at 4 °C. The supernatant was removed and the pellets were each divided into 500 μl LB medium. Lambda phage solution (1.21·10^9^ pfu/ml) was added to the bacteria at dilutions of 1:50, 1:500 and 1:5000. The phages were allowed to adsorb for 1 min. After adsorption, free phage particles were removed by centrifugation. The pellet was resuspended in 1 ml LB medium. The preparations were each added to 99 ml LB medium and incubated at RT and 120 rpm. At 5 min intervals, 250 μl samples were taken, mixed with 5 ml of soft agar [0.3% (w/v)] and poured over an LB plate. Resulting plaques were quantified following over-night incubation at 30°C. The burst size was calculated by dividing the average phage titer at the plateau phase by the average phage titer along the latent phase.

### Membrane depolarization assay

Bis-(1,3-dibutylbarbituric acid) trimethine oxonol (DiBAC_4_(3); Molecular Probes^TM^) was used for staining of depolarized cells. A 10 mg/ml (w/v) stock solution was prepared in DMSO. Working solutions [200mg/ml (w/v) in water and 0,5 % Tween20] were prepared freshly. About 500ml liquid culture aliquots were collected by centrifugation (13,000 rpm, 3 min). All steps were performed at room temperature. Pellets were resuspended in 200 ml PBS and stained with DiBAC_4_(3) at a final concentration of 1mg/ml (w/v) for 20 min in the dark. After three steps of washing with 200 ml PBS, cells were resuspended in 200 ml PBS and evaluated using the TECAN plate reader. The used bis-oxonal has an excitation maximum of 490 nm and an emission maximum of 516 nm.

### Bioinformatic approaches

The gene map of the LambdaSO genome was generated using the SnapGene 7.0 software (snapgene.com). The primary sequence of the potential spanin proteins was characterized regarding a prediction of protein domains using SMART (46), SignalP (47) and DeepCoil (48). Topological prediction and classification of transmembrane proteins was carried out using the DeepTMHMM database (49). ProtParam (50) was used for the calculation of physical and chemical parameters of proteins. The structure of all lysis-involved proteins of LambdaSo was predicted using Alphafold2 (51). The obtained data was then analyzed using Pymol (The PyMOL Molecular Graphics System, Version 2.0 Schrödinger, LLC) and the included APBS electrostatics tool. The NCBI and Pfam database were accessed to obtain the aminoacid sequences of other phage lysis proteins.

## Acknowledgements

The authors would like to acknowledge Ulrike Ruppert for excellent technical support. The work was supported by a grant (TH831/9-1) from the Deutsche Forschungsgemeinschaft (DFG) within the priority program SPP2330.

